# High-throughput multicolor 3D localization in live cells by depth-encoding imaging flow cytometry

**DOI:** 10.1101/730101

**Authors:** Lucien E. Weiss, Yael Shalev Ezra, Sarah E. Goldberg, Boris Ferdman, Yoav Shechtman

## Abstract

Imaging flow cytometry replaces the canonical point-source detector of flow cytometry with a camera, unveiling subsample details in 2D images while maintaining high-throughput. Here we show that the technique is inherently compatible with 3D localization microscopy by point-spread-function engineering, namely the encoding of emitter depth in the emission pattern captured by a camera. By exploiting the laminar-flow profile in microfluidics, 3D positions can be extracted from cells or other objects of interest by calibrating the depth-dependent response of the imaging system using fluorescent microspheres mixed with the sample buffer. We demonstrate this approach for measuring fluorescently-labeled DNA *in vitro* and the chromosomal compaction state in large populations of live cells, collecting thousands of samples each minute. Furthermore, our approach is fully compatible with existing commercial apparatus, and can extend the imaging volume of the device, enabling faster flowrates thereby increasing throughput.

## INTRODUCTION

Flow cytometry enables the rapid characterization of large populations of cells and particles. Increasing reliability and decreasing costs have helped the technology emerge at the forefront of medical diagnostics, drug-efficacy screens, cell sorting and characterization, personalized medicine, and more (Betters, 2015; Brown and Wittwer, 2000; Luider et al., 2004). While fluorescence intensity, side/forward scatter, and absorbance measurements are now standard capabilities of modern, commercially-available instruments, subsample characterization is typically not possible when the signal is relayed onto point detectors, thus the interpretability of acquired data is constrained to relative measurements and signature recognition by comparison to standards (Chandler et al., 2011).

Imaging flow cytometry (IFC) replaces the canonical point detector(s) with a 2D camera, thus enabling widefield microscopy at high throughput (Cambier et al., 1979; Kay and Wheeless, 1976; Kay et al., 1979). Recently-developed instruments have demonstrated multiple spectral channels (Basiji et al., 2007; George et al., 2004) and other microscope modalities, *e.g*. phase imaging (Gorthi and Schonbrun, 2012) and optical sectioning (Gualda et al., 2017; Regmi et al., 2014). In general, because the basic operating principles of IFC and widefield microscopy are the same, tools developed for one can be applied to the other.

One widefield technique that has been applied to attain nanoscale information is localization microscopy (Toprak and Selvin, 2007). The central premise is that spreading out the emission light of a point source over several camera pixels enables the underlying emitter’s position to be extracted with a precision far below the pixel size. This approach has two main requirements: 1) a sufficiently-low emitter density in each image such that individual emitters can be recognized. 2) Knowledge of the optical system’s point-spread function (PSF), namely what the expected distribution of light emanating from a point-source is on the detector.

The low-emitter-density requirement can be satisfied, to some extent, by flow cytometry, where the microfluidics are designed to pass objects through the imaging region one at a time. Knowledge of the PSF is used to extract the underlying parameters from an image, namely an emitter’s likely position. Note that dense yet small volumes of emitters, such as small fluorescent beads or punctate spots in cells, can be treated collectively as single point sources, as long as they occupy a sub-diffraction volume.

For 2D localization microscopy, the in-focus PSF is often approximated by a symmetric, 2D-Gaussian function, which is used to fit the measured image. This model can account for some defocus, which manifests as broadening (or blur) in the object’s image; however, the approach is poorly suited for 3D microscopy because there is a redundant widening in the PSF as the object is displaced in-front-of or behind the focal plane (Kao and Verkman, 1994). This ambiguity is resolved in iPALM and biplane microscopy by imaging on multiple detectors (Abrahamsson et al., 2012; Juette et al., 2008; Shtengel et al., 2009). Alternatively, by adding an aberration into the imaging system *via* insertion of additional optical elements, the PSF itself can be modified to remove the ambiguity. This technique is broadly defined as PSF-engineering microscopy and is advantageous in that it can be implemented in a standard, single-camera imaging system (Backer and Moerner, 2014).

A simple implementation of PSF engineering is achieved by adding a cylindrical lens into the imaging path, encoding the depth into the directional stretching of the PSF shape (Huang et al., 2008; Kao and Verkman, 1994). The result can be fit with an asymmetric 2D-Gaussian function, then processed *via* a shape-to-depth calibration curve, which is normally generated by imaging a stationary object at various defocus levels (Huang et al., 2008; Mlodzianoski et al., 2009). More complicated PSFs, realized by adding a specially-designed, phase-shaping optical element into the microscope’s back focal plane (BFP), have been used to extract 3D spatial information (Baddeley et al., 2011; Pavani et al., 2009; Shechtman et al., 2014) as well as to directly encode additional emitter information into an image, *e.g*. dipole orientation (Backer et al., 2013; Backlund et al., 2012) and emission wavelength (Shechtman et al., 2016; Smith et al., 2016). In a related yet disparate application, the incorporation of diffractive optical elements for enhanced-depth-of-field (EDOF) (Dowski and Cathey, 1995) has been applied to IFC, modifying the PSF of the microscope to make it depth independent (Ortyn et al., 2007), which effectively produces a 2D projection of the imaged volume.

Here, we demonstrate high-throughput, nanoscale, 3D multicolor localization by combining IFC with PSF engineering. Furthermore, we show how this method can be implemented directly into existing instruments with a minor, removable addition. The method is comprised of the following four steps: 1. inserting an optical element (*i.e*. a phase mask or a cylindrical lens) into the imaging path of an imaging flow cytometer; 2. capturing images of flowing objects (*e.g*. fluorescently-labeled cells) along with a calibration sample, i.e. fluorescent beads, that follow a known (or measureable) depth distribution function; 3. Generating a calibration curve by decoding the PSF response for the calibration sample; and 4.applying the calibration curve to obtain the 3D positions of the sample images, which can then be colocalized between color channels.

We first validate the approach by obtaining large 3D datasets of fluorescently-labeled DNA in *vitro*: by imaging DNA-origami nanorulers, and *in vivo*: by measuring chromosomal compaction states inside live yeast cells. Lastly, we show how PSF engineering can extend the depth range in IFC using the Tetrapod PSF (Shechtman et al., 2015). Importantly, the flow-based experiments presented here were performed using a commercially available, multicolor, imaging flow cytometer, the Amnis ImageStreamX (ISX) with the supplied acquisition software. The output of the IFC software was processed with custom MATLAB scripts, thereby demonstrating the immediate applicability of our approach to existing apparatus.

## RESULTS

### Device and functionality

An illustration of the device is shown in Figure 1 A. Samples are loaded into the instrument and then directed through a multilaser-illuminated imaging volume. Emission light from the sample is captured by a 60X objective, relayed through a series of lenses, divided into separate spectral channels, and directed onto a camera, whose readout rate is synchronized with the flow speed. As a first demonstration, we show how an astigmatism can be incorporated into the system. By inserting a cylindrical lens between two of the aforementioned relay lenses (see Methods), the axial and lateral focal planes of the instrument are bifurcated, thus an object closer to the objective will appear focused laterally, but defocused axially and *vice versa* (Figures 1 B, C). The extent of the astigmatism contains information about the depth of the emitter, but is not a direct measurement of the z position. The typical 3D calibration used in microscopy (Huang et al., 2008) is therefore inapplicable and the incorporation of PSF engineering into flow imaging necessitates a novel 3D calibration procedure, which we describe below.

**Figure 1.**
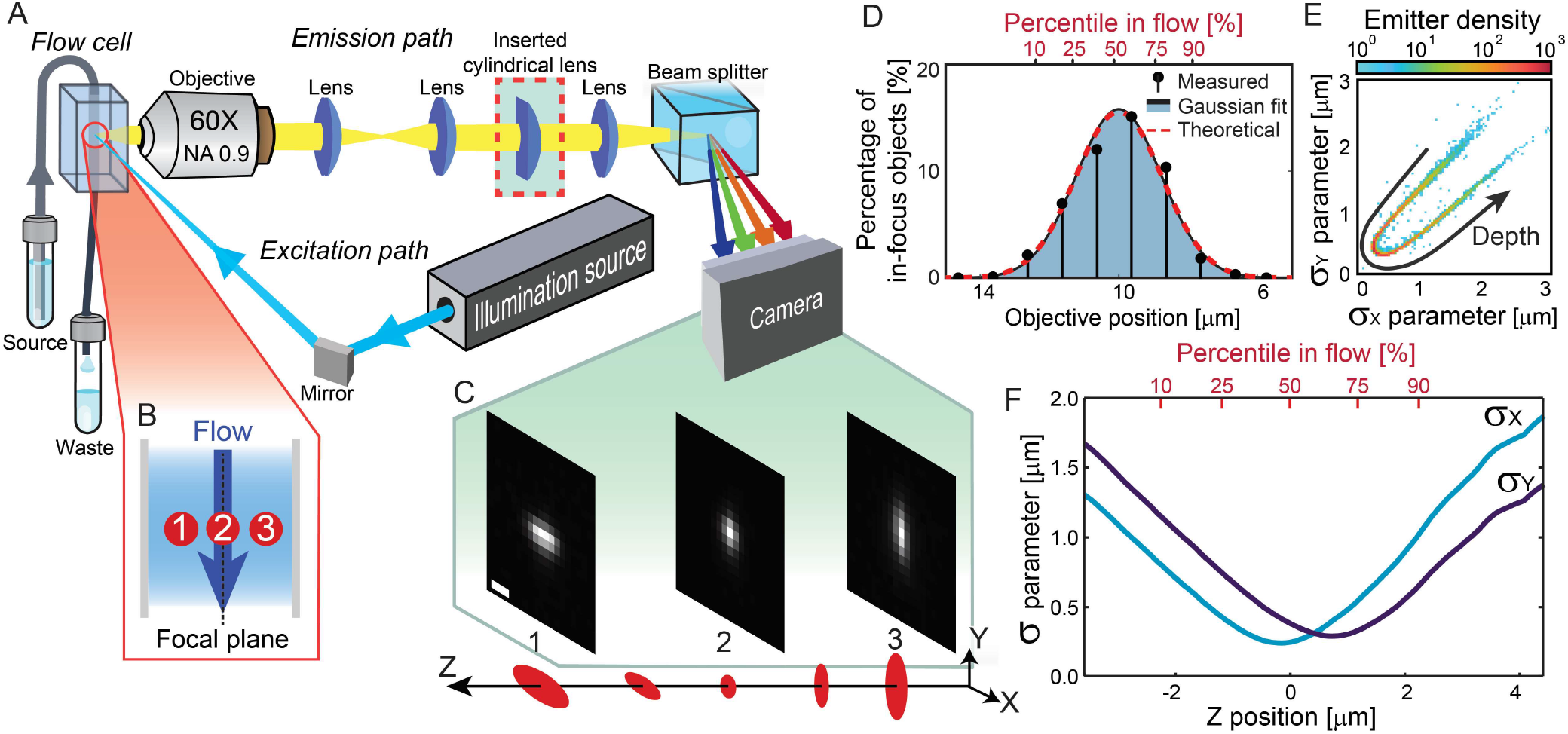
Schematic of the apparatus and 3D calibration. (A) Objects directed through a flow cell are illuminated with a laser light source. The emission is collected by a 60X air objective, and the light is directed through a tube lens, then two relay lenses with a cylindrical lens inserted between them (red box), before being split into spectral channels and imaged by a detector. Not to scale. Objects at different depths in the flow channel (B) have a (C) depth-dependent point-spread function imaged on the detector, scalebar 1 μm. (D) The distribution of objects measured in the flow. (E) The distribution of localized object astigmatisms, where the density is given in *emitters per 10 nm^2^* in terms of the Gaussian shape parameters σ_X_ and σ_Y_. (F) The shape-to-depth calibration curve generated from (D) and (E).

Unlike a standard microscope, whose PSF can be calibrated by z-scanning a stationary fiducial (*e.g*. a fluorescent bead), in a quickly-moving flow, it is not possible to scan a point source to characterize the PSF as a function of z. Our solution for obtaining an accurate experimental-PSF calibration is to exploit the statistical properties of the flowing beads. In brief, since the probability distribution of object depths (z positions) in laminar flow is well defined and measureable (Figures 1 D), a sufficiently-large sample size will closely resemble the average distribution. A dataset of calibration images can therefore be obtained and algorithmically sorted by relative depth, *e.g*. by the extent of each image’s measured astigmatism to produce a shape-distribution curve (Figure 1 E). This astigmatism curve is then parameterized by the depth percentile, where each point along the curve, *i.e*, shape-parameter pair is assigned the statistically most-likely z position (Figure 1 F). As the number of calibration measurements increase, the average position error decreases (Figure S1). New images can then be analyzed and compared to the reference curve to find their depth irrespective of their probability distribution in the flow, namely, after calibration, any point-like object can be localized in 3D.

### Six-color colocalization of fluorescent beads

While determining the 3D positions in a single color channel can be used to measure the flow profile and characterize the PSF (Cabriel et al., 2018; Shechtman et al., 2015), of more practical interest in imaging cytometry is measuring the relation between two or more objects. For emitters separated by distances below the diffraction limit (^~^300 nm), this can be accomplished by splitting the emission of the two or more objects onto different images, *e.g*. by spectrally separating the objects. To assess our ability to utilize the six spectral channels collected by the ISX, and characterize our method’s performance, we employed 200 nm Tetraspeck fluorescent beads due to their broad emission spectrum (see Methods). We collected and analyzed a dataset of 50,000 beads s, each with six images in less than 3 minute. Next, we divided the data into two independent groups: one for calibration and one for evaluating the performance of our approach, namely, the colocalization precision between spectral channels. Images of an example bead are shown in Figure 2 A, where the localized positions, relative to the mean, are shown in Figures 2 B & C. The inter-channel residual distances are plotted as 2D histograms in Figure 2 D, and were used to compute the geometric 3D distances per image. The cross-channel error was defined as the median 3D distance (Figure 2 E), and ranged from 25-to-53 nm for various channel pairs in optimized conditions (*i.e*. cylindrical lens placement, laser power, and flow stability).

**Figure 2.**
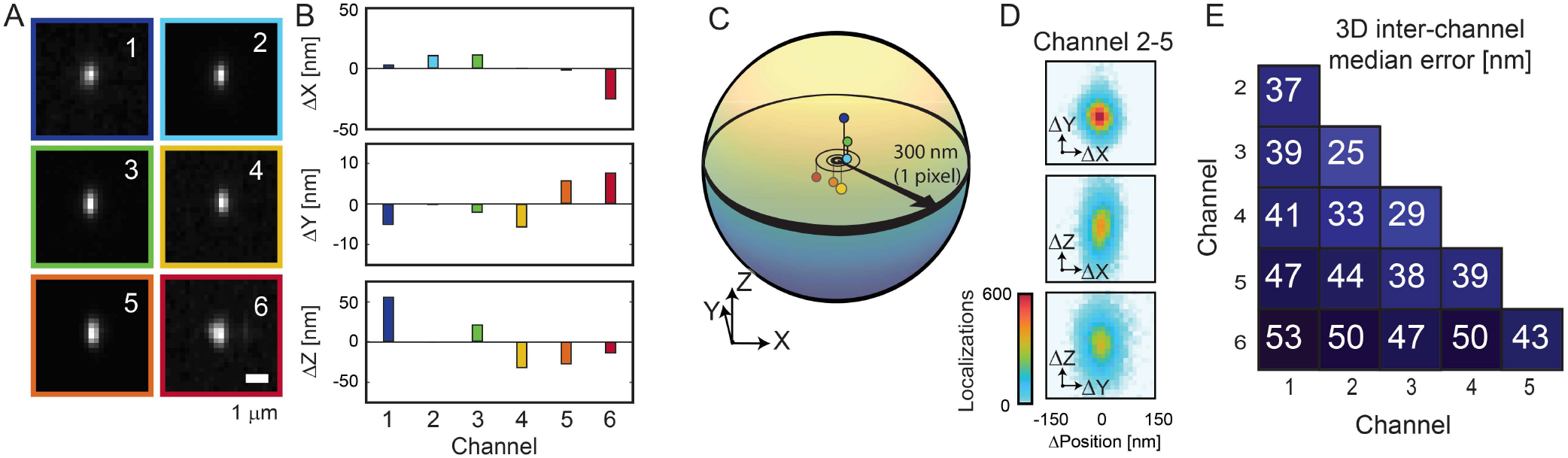
Six-color, 3D colocalization. (A) Six measurements of the same fluorescent bead recorded in different spectral channels. (B) The localized positions in each color channel relative to the mean. (C) The 3D positions of the localizations in each color channel. (D) 2D histograms showing the difference in localization position for 25,000 beads. (E) The median 3D distance for the 25,000 beads between different channel combinations.

### 3D distance measurements of DNA nanorods

To demonstrate 3D distance measurements between pairs of objects, we employed a DNA-origami structured nanoruler where the two ends were labeled with green (Atto488) and red (Atto647N) dyes, with a designed separation of 180 nm (Schmied et al., 2014), (Figures 3 A & B). The sample was diluted and mixed with fluorescent beads prior to imaging. A subset of fluorescent beads was used for calibration and the remainder for comparison to the nanorulers. The calculated inter-channel distances in each axis (Figure 3 D) was used to compute the geometric 3D distance between the probes per object, *i.e*. to calculate the object length. We found the results obtained in flow, 164 ± 87 nm (mean ± standard deviation) (Figure 3 E), was similar to that observed by conventional 2D microscopy where the nanorulers were fixed to a glass coverslip, 151 ± 66 (mean ± standard deviation) (Figure S2).

**Figure 3.**
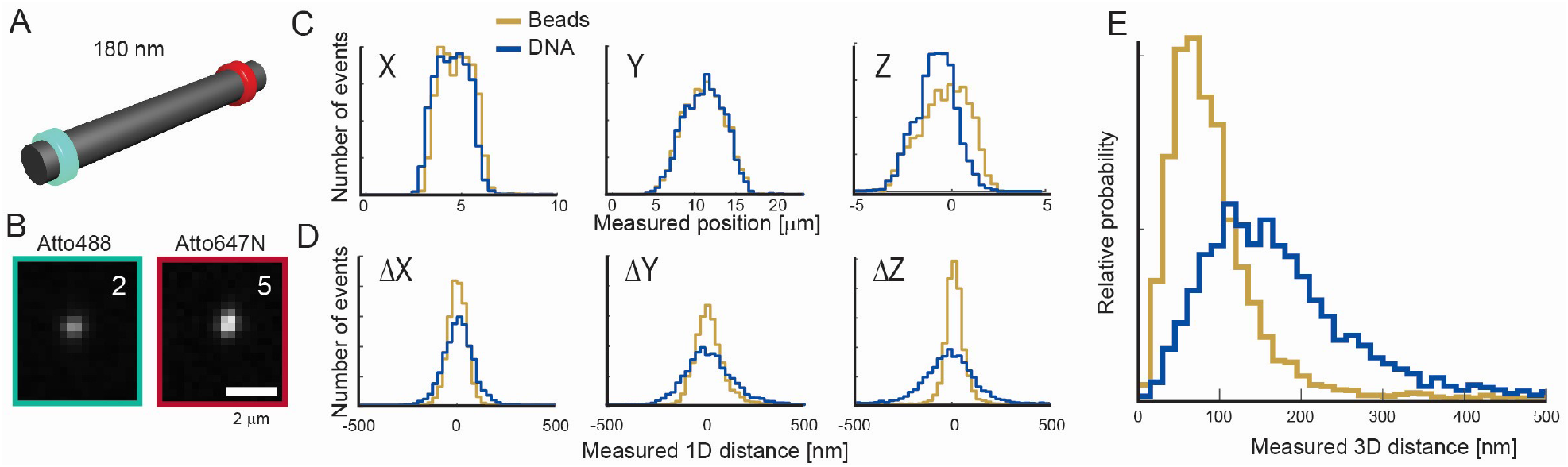
3D measurements of DNA nanorulers. (A) An illustration of a DNA origami-based nanoruler, where the two ends are labeled with Atto488 and Atto647N fluorophores. (B) An image of a single nanoruler imaged in two channels. (C) A histogram of localized positions in the two color channels. (D) A histogram of the inter-channel distance between localizations in each dimension. (E) A histogram of the 3D distance measured. Standard microscopy images and analysis are shown in Figure S2.

### Measuring chromosomal compaction states in Saccharomyces cerevisiae

Populations of live cells show temporally-dependent and highly-variable responses to stimuli which necessitates high-throughput imaging. For example, chromosomal repair after ultraviolet-induced DNA damage (Aylon and Kupiec, 2004), responses to changes in the chemical environment (Dultz et al., 2018), and gene position through the cellular cycle (Shachar et al., 2015), and gene regulation (Khanna et al., 2019) have all been shown to involve dynamic reconfiguration of the chromosome in 3D.

To demonstrate the applicability of our approach to these types of biological measurements, we imaged fluorescently-tagged DNA loci of live yeast cells to investigate a proposed mechanism for gene regulation: regions of the chromosome containing suppressed genes are spatially compacted (Even-Faitelson et al., 2016; Gilbert et al., 2004; Wolffe and Pruss, 1996). In brief, two arrays of operator sites were integrated into the chromosome flanking the Gal locus (Dultz et al., 2018), a region encoding several genes for the galactose-metabolism machinery (Hopper et al., 1978). Fluorescently-labeled operator-binding proteins attach to each operator array forming two fluorescent spots (Figure 4 A), where the inter-spot distance is affected by the growth conditions (Figure 4 B).

**Figure 4.**
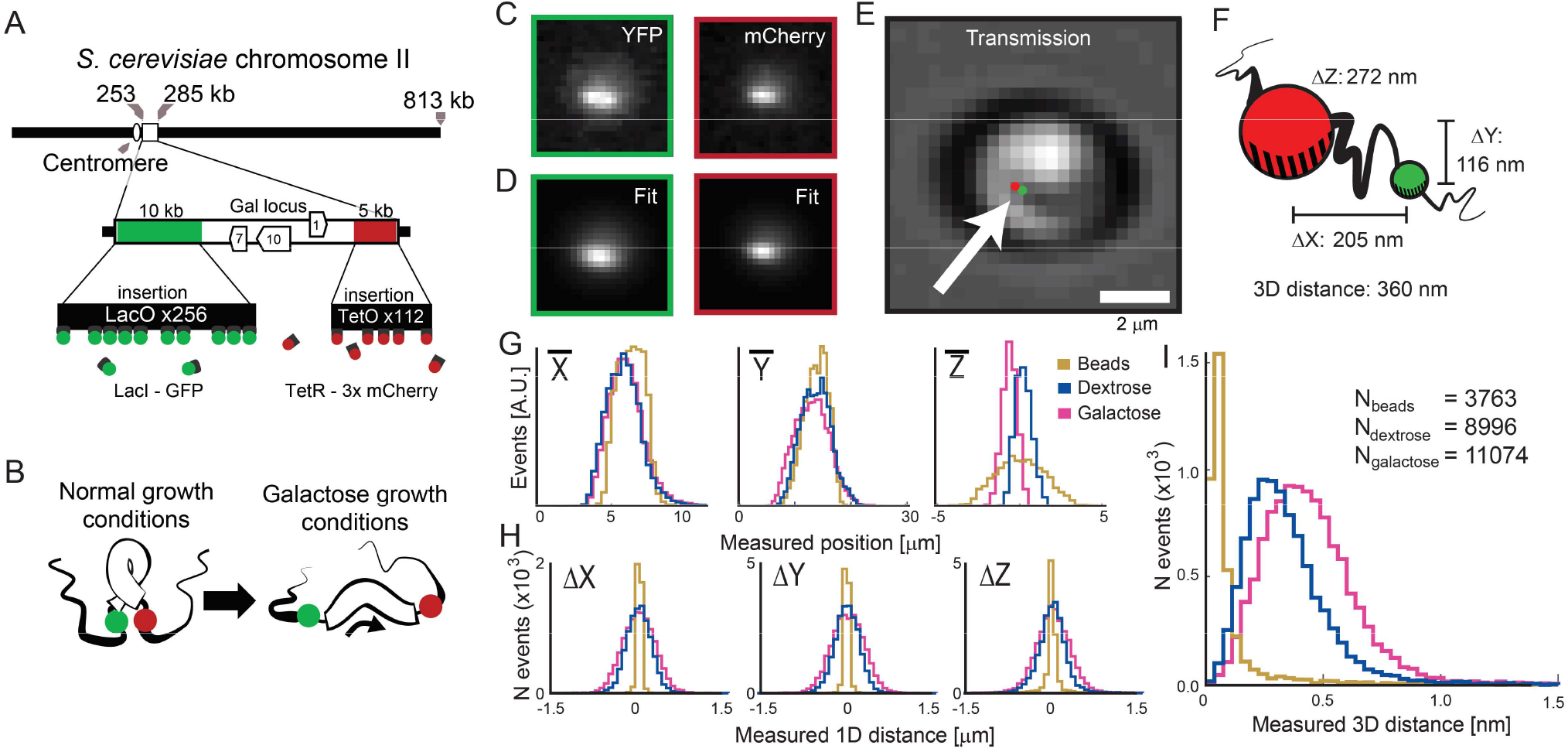
3D compaction of the Gal locus in live cells. (A) Schematic of the fluorescent-labeling scheme. (B) Illustration of the chromosome compaction states. (C) Images of live yeast cells in fluorescence with corresponding (D) model images resulting from the localization. (E) Transmission image with the two localized positions superimposed, highlighted by the white arrow. (F) The 3D displacement measured from the same cell. The black line connecting the two points is meant to illustrate the flexible DNA linker (G) Histograms of the localized positions in each axis. (H) Histograms of the measured displacements between channels in each axis and (I) 3D. The results obtained for fixed cells are shown in Figure S3.

To account for the cellular autofluorescence and unbound fluorescent proteins, our localization algorithm was modified to include an additional Gaussian-background term (Figure 4 C & D). Each cell image was then analyzed in the two color channels separately to obtain absolute 3D positions (Figure 4 E & G) which were then used to calculate the relative displacement in each dimension separately and 3D (Figure 4 F H & I). Similar to previously measurements (Dultz et al., 2016; Shechtman et al., 2017), the mean inter-loci distance was observed to be dependent on the growth condition (mean ± standard deviation: 339 ± 174 in dextrose and 434 ± 193 in galactose). Crucially, while previous 3D datasets were recorded over hours at ^~^1 image per second (Shechtman et al., 2017), the current method acquires hundreds of images per second, producing a large library of suitable cell images in only a few minutes (N = 8,996 for dextrose and 11,074 for galactose, recorded in <5 and <10 minutes, respectively). A similar change in the inter-loci distance was observable for cells that were fixed prior to imaging (Figure S3, Methods). To estimate the 3D colocalization precision, we colocalized the same position on the chromosome marked with mCherry in two spectral channels (4 & 5), which was possible due to the broad emission spectrum of the fluorophore. The 3D colocalization precision was found to be 160 nm ± 100 nm (mean ± standard deviation).

### Extended imaging depth using the Tetrapod point-spread function

To demonstrate the generality of our approach to other point-spread functions, we inserted another phase-shaping optical element into the imaging path, replacing the cylindrical lens (Figure 5 A). The Tetrapod phase mask was designed to optimize the Fisher information for 3D localization (Shechtman et al., 2014), and can be used for 3D measurements over extended depths (Figure 5 C). While in the previous measurements, we used the extent of the astigmatism to order images by depth, this solution is not applicable to any 3D PSF, namely, those that cannot be easily fit with an astigmatic Gaussian. Here, we employed a more general solution to generate the calibration curve with no assumed knowledge of the PSF. First we collected 23 datasets of 5,000 beads flowing with a narrow-depth distribution (5 μm), where the objective lens was moved forward by 2 microns between each dataset. The images within each dataset were aligned by a center of mass calculation based on their intensities and averaged to produce an image template. Next, a representative image was identified in each dataset by calculating the 2D correlation of each image with the template. These selected images composed an intermediate calibration dataset, as they were already ordered by z. We then increased the flow diameter to 50 μm (Figure 5 B) and imaged 50,000 PSFs, ranking them by their similarity to the images in the calibration dataset. These ranked images were then assigned a probable depth from the known flow dimensions (Figure 5 D) thus creating our shape-to-depth calibration. As an additional verification procedure, the experimental results were compared to a numerical simulation of the optical system (Figure 5 E, see Methods).

**Figure 5.**
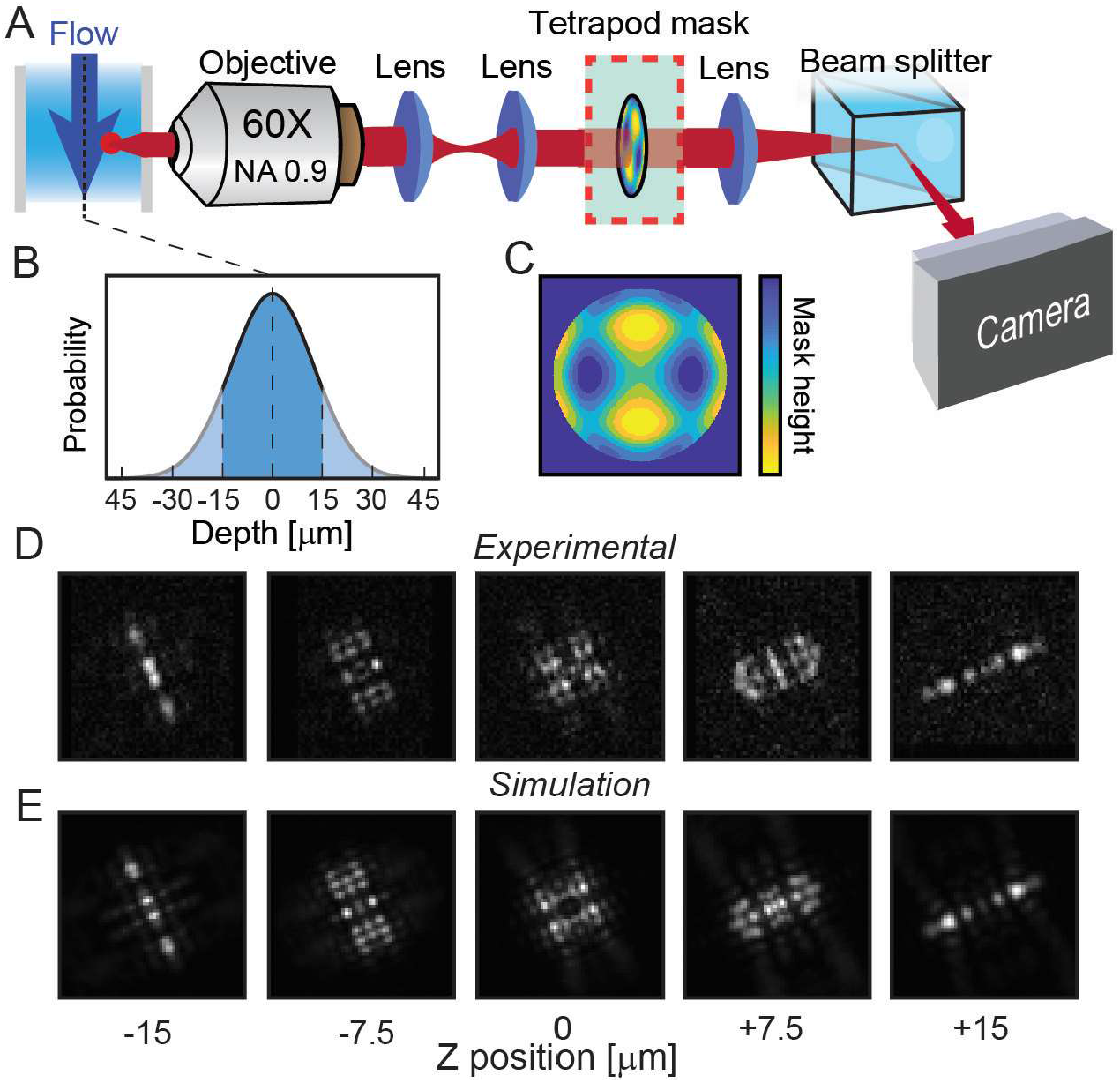
Extended depth-range 3D localization with the Tetrapod point-spread function. (A) A schematic of the emission path of the imaging flow cytometer. (B) The extended depth distribution imaged. (C) A close-up of the Tetrapod phasemask pattern. (D) Experimentally measured PSFs within the instrument. (E) Simulated images at depths corresponding to the calculated depths of objects shown in (D).

## Discussion

In this work, we have extended the capabilities of imaging flow cytometry to enable three-dimensional localization. This was made possible by taking advantage of the high-throughput nature of IFC to derive a 3D model of the PSF using the underlying flow distribution. Furthermore, we have shown that our approach is fully compatible with existing commercial apparatus and requires only a small hardware addition to the device: the placement of a cylindrical lens or phase mask into the imaging path. Using this approach, we have increased the throughput of 3D-distance measurements of DNA compaction in live yeast cells by orders of magnitude over previously utilized traditional-microscopy methods (Shechtman et al., 2017).

In our measurements, we identified three sources of error: field-dependent aberrations, finite localization precision (both occur in any type of localization microscopy), and finally, the synchronization of the flow rate with the camera acquisition (unique to IFC). Field-dependent aberrations come from imperfections in the optical system, *e.g*. misalignment of the cylindrical lens. To account for systematic biases as a function of the emitter position within the flow, localizations were adjusted using an interpolated 3D correction map generated from the calibration beads (Figure S4).

To characterize the localization precision as a function of signal level, we measured beads under different illumination intensities and colocalized them in two channels (2 & 4). As expected, higher signal intensities translated to improved precision (Figure S5); however, at sufficiently high signal intensities (a mean amplitude >50 D.U., as was the case for the DNA nanorulers), another source of error often dominated the attainable 3D precision. The source of this error is desynchronization of the acquisition rate with the flow rate, which causes images to be slightly blurred in the direction of flow (Y). Future improvements that stabilize or produce more accurate determinations of the flowrate will reduce this error.

Like all localization-based methods, a suitably low density of emitters is required so that the PSFs do not significantly overlap in the image. While certain types of objects, such as fluorescent beads, are ideal for this application, we made use of the multiple spectral windows imaged by the ISX to colocalize closely spaced regions of DNA on fluorescent nanorulers and selectively-labeled regions of the chromosome in live cells, where the close proximity of the imaged spots would be overlapping in a single-channel wide-field measurement. Using spectral separation of fluorescence emission, however, is not a fundamental requirement of this or other localization microscopy methods. The recent application of machine learning, namely neural networks (NNs), to localization microscopy has been shown by our group (Nehme et al., 2018, 2019) and others (Boyd et al., 2018; Ouyang et al., 2018) to be adept at extracting the positions of multiple closely-spaced emitters (>6 emitters/μm^2^), and could represent one path to enable more complex samples. Favorably, NNs also have the advantage of rapid data processing, and have recently been applied to online analysis controlling cell sorting during IFC experiments in a custom instrument (Nitta et al., 2018).

In addition to new applications directly related to IFC, the approach described here may be used to calibrate any microscope’s 3D PSF by utilizing a flow system with well-known depth-distribution properties to produce a calibration curve that could be applied to static samples. This approach could potentially solve the long-standing challenge caused by using calibration curves generated with surface-adhered objects, when imaging is subsequently performed in media with a different refractive index (Egner and Hell, 2006).

Related to IFC, the work presented here does not exhaust the full potential of combining PSF engineering to flow imaging. Future applications could use customized phase masks design to make optimal use of the flow profile (z-positions) in a particular experiment (Shechtman et al., 2015), and make the depth determination more robust to variations in flowrates, *e.g*. by encoding the depth in a PSF stretch in one direction (Backer et al., 2014; Jia et al., 2014) orthogonal to the flow, decoupling the depth-estimation error from flow-rate induced errors. PSF engineering could also be used to enhance the autofocusing and core-size characterization by applying PSF engineering to a dedicated imaging channel inside an instrument. Finally, it would be useful to incorporate fluorescence-activated cell sorting (FACS) based on sub-cellular 3D colocalizations by incorporating online analysis and classification of cells into the IFC operating software, which provides intriguing possibilities for new types of selection-marker technologies.

## Acknowledgments

We thank E. Barak and Y. Lupu-Haber and the Lorry I. Lokey Interdisciplinary Center for Life Sciences and Engineering for technical assistance with the imaging flow-cytometer. The multicolor yeast cells were provided by K. Weis and E. Dultz. The Tetrapod PSF was fabricated by M. Y. Lee. This work was supported by the European Research Council (ERC) under the European Union Horizon 2020 research and innovation program (Grant no 802567), the Zuckerman foundation, and the POLAK Fund for Applied Research, at the Technion.

## AUTHOR CONTRIBUTIONS

LEW & YS conceived of the approach. LEW, YSE & SEG prepared samples and conducted the experiments. All authors analyzed the data and contributed to the manuscript.

## METHODS

### Sample preparation for flow cytometry

Prior to loading into the ISX, samples were diluted to < 2 × 10^5^ objects per μl and mixed with fluorescent beads. This density was chosen to balance a high acquisition rate (≥ 100 objects per second) while keeping the probability of imaging multiple particles at the same time relatively low. Fields of view containing two or more objects were either cropped or discarded.

For calibration, 0.2 μm diameter TetraSpeck (TS) beads (Invitrogen, cat. T7280) were diluted 1:200 in 1X PBS (phosphate-buffered saline, Sigma cat. P5368 dissolved in 1L ddH_2_O; NaCl 0.138 M, KCl 0.0027 M, pH 7.4) to a final density of 1 × 10^5^ beads per μl, which translated into ~150 objects imaged per second.

DNA nanorulers (GATTAquant GmbH) with a designed length of 180 nm separating two groups of emitters at either end (ATTO 647N and ATTO 488) were prepared for IFC as follows: 2-4 nM stock was diluted 1:200 in 1xTAE/10mM MgCl_2_. (1xTAE contains 4.844 g Tris, 1.21 ml acetic acid, and 0.372 g EDTA in 1L ddH_2_O). Fluorescent beads were then added as described previously.

Yeast cells were cultured by standard growth protocols (Bergman et al., 2001) at 30°C, 200 rpm. In brief, cultures were chosen from single colonies grown on non-selective YEPD agar. Cells were grown overnight in synthetic depleted low-fluorescence media (-Tryptophan) (SD(-Trp)-LFM), (Formedium, cat. CYN6501) and added amino acids, along with 2% (w/v) raffinose (Alpha Aesar, cat. A18313). Cultures were then diluted in synthetic complete low-fluorescence media (SC-LFM) and either 2% galactose (Acros, cat. 150615000) or 2% (w/v) dextrose (Sigma, cat. G5767). After three more hours of incubation, cells were imaged in the mid-log phase 0.5<O.D_600_<1 (~0.7), without centrifuging. For 3D calibration, fluorescent beads were added immediately before imaging to match the number of cells imaged per second.

All buffers and media used for cell growth and imaging was sterilized by filtering prior to use (0.22 um).

### Inserting the cylindrical lens and phase mask into the Imaging Flow Cytometer

A cylindrical lens (*f* = 1 m, Thorlabs, LJ1516RM-A) was mounted in a 2D micrometer translational mount (Thorlabs, LM1XY) and inserted into the imaging path of the ISX using a detachable magnetic base (Thorlabs, KB2X2). The position of the lens was then adjusted until the astigmatic PSFs appeared symmetrical and oriented vertically/horizontally across all channels. The position along the optical axis changes the nature of the astigmatism shape, and was optimized iteratively.

The Tetrapod phase mask was mounted in a custom adapter designed to fit in the filter wheel located at a plane conjugate to the back focal plane of the objective. The mask position was refined in X and Y until the PSF was symmetrical.

### Imaging flow cytometry settings

The default ISX parameters were used for data collection with the exceptions described below. The ISX was used with the 60X magnification objective and a Core Diameter 7 μm (default for 60X magnification) except when specified otherwise, namely, in the Tetrapod experiment. Autofocus was set to OFF (default was ON), and the ideal focal position was found after inserting the cylindrical lens. This was done by imaging short acquisitions ^~^2 minutes of fluorescent beads and minimizing the distance between the average focal plane (where the shape of the object is symmetric) and the center of the flow, that is to say the number of objects on either side of the focal plane was approximately even. This value was typically in the range of [-2.5, 2.5] μm and appeared robust between measurements on the same day. Next, after enabling the Advanced Configuration setting, ObjectMaskCoeffParam was set to 0 (default was 0.8), and Keep Clipped Objects was set to ON (default was OFF) to retain the largest field of view possible; note that these two settings were only needed for the Tetrapod PSF, but were used in all measurements described here for consistency.

For beads and cell data, the sheath buffer was 1X PBS; however, for the DNA nanoruler experiment, the sheath buffer was exchanged with 1X TAE/10mM MgCl_2_ to avoid possible changes in origami structure due to changes in ionic strength.

### Imaging flow cytometry data preparation for 3D analysis

There were two steps of data preparation before 3D localization: object classification, and assignment of offset X, Y position.

Bead and non-bead objects were classified using the data in the exported feature file generated by the IDEAS companion software supplied with the ISX. The feature *Image_MC<channel number>* provided sufficient classification. According to the IDEAS manual, this feature is “the sum of the pixel intensities in the mask, background subtracted”. The described classification scheme worked well due to the differences in fluorescence intensity and ratio between some spectral channels of the fluorescent beads relative to both the cells and the nanorulers. More general classification schemes might require use of the full image data rather than relying on exported feature parameters. Note that in the case of highly similar calibration and test samples, it may be necessary to measure the PSF calibration separately. Note that we did observe some subtle differences in the PSF calibration once the flow was stopped and then restarted, likely due to a change in the objective position which resets upon changing samples. It is therefore best to collect calibration data while imaging the sample.

The absolute XY position of an emitter with respect to the relevant CCD channel is the sum of its position in the image frame and the offset of the image frame with respect to the CCD channel. The offsets in X and Y are the exported feature parameters *Raw_Centroid_X* and *Raw_Centroid_Y*. Note that retaining the absolute X and Y position for each object is necessary to correct for any systematic bias that occurs while fitting the astigmatic PSF when the cylindrical lens is not perfectly aligned.

### Software

Imaging flow cytometery datasets (.cif files) were generated using INSPIRE software (Amnis). Post-experiment, feature data for all objects was exported to .txt format using IDEAS software (Amnis). The feature data was useful for rapid classification of objects as either calibration beads or cells/DNA nanorulers.

Matlab (Mathworks, version 2018b) with Bioformats package (Linkert et al., 2010) was used for analysis of all .cif data from cell and nanoruler samples.

### Localization of objects

In all data acquired with the cylindrical lens, intensity images ***I***(***x, y***) of fluorescent objects were fit using nonlinear least-squares (Matlab’s *lsqnonlin* function) to the following 2D Gaussian model:

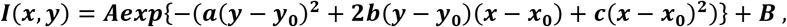

where ***B*** is a constant background intensity for bead and *in vitro* DNA data, ***A*** is the maximum intensity, and (***x*_0_**, ***y*_0_**) is the x-y position of the emitter. The widths, 2***σ*_1_** and **2*σ*_2_**, of the 2D Gaussian and its rotation θ from the CCD axis are related to ***a, b*** and ***c*** as follows:

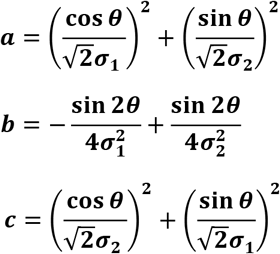

We defined the axes of the 2D Gaussian as follows: the positive ***σ*_2_** axis is rotated 90º clockwise from the positive ***σ*_1_** axis. The positive axis of ***σ*_1_** may be rotated in the range of (−45º, 45º) with respect to the positive Y axis of the CCD. This results in the positive axis of ***σ*_1_** aligned closer to the Y axis of the CCD, and the positive axis of ***σ*_2_** aligned closer to the X axis of the CCD.

Emitters within cells were fit in the same way, except that the background was also a 2D Gaussian function, whose shape parameters were restricted to being larger than that of the localized emitters inside the cell:

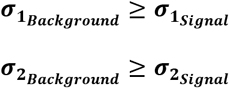

### Parameterizing the Gaussian fits and localizing in 3D

To parametrize the PSF shape, namely (***σ*_1_**, ***σ*_2_**) as a function of the emitter depth, one or more channels was used for calibration. After initially fitting with a freely-rotating astigmatic Gaussian, the precise orientation angles of the Gaussian were found per channel by taking the mean angle of calibration beads whose shape parameters were between [1, 3.5] pixels with a finite ratio [1.1, 2] pixels. The average angle for each side of focus was determined separately to account for aberrations in the imaging system that distort the PSF. Images were next re-localized with the fixed angles identified.

Next, to create a preliminary ordering of the calibration objects images by depth, a 16-parameter function was used to generate a 2D image to match the 2D histogram of shape distributions (Supplemental code will be made available online), Figure 1 E. Once a curve that approximated the range of object shapes was found, the data was projected to the closet point along the curve to find the relative order of the objects. If more than one channel was used for calibration, the average object position was used. Next, in each channel analyzed in 3D, objects were binned by their relative depth position to find the average shape parameters for percentiles of data in each channel. Finally, a spline function to create a continuous function for the shape as a function of percent along the flow (0-100%), Figure 1 F. To convert the percentile in the flow distribution to an absolute Z position, we utilized the Matlab function *norminv* which outputs a position as a function of percentile for a normal distribution of a given mean (in our case 0) and standard deviation (^~^1/4 of the flow diameter). All images were then analyzed as a function of X, Y, and Z.

### Validating the depth distribution within the stream

We determined the distribution of particle density as a function of position within the core of the flow as follows: we first collected data from fluorescent beads for different positions of the focal plane with respect to the core in 1 μm increments, over a range of 10 μm, with constant core diameter of either 5 or 7 μm and in the absence of the astigmatic lens. The fraction of objects in focus at each distance from the objective ***f*** was defined as:

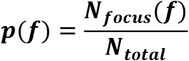

where ***N_total_*** is the total number of objects collected, and ***N_focus_*** is the number of objects for which 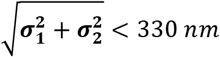 (1.1 pixels). The in-focus threshold was chosen to be small enough so that **Σ*_f_p_f_*** <1, indicating that objects were not defined to be in focus for more than one focal position ***f***, and large enough to include a large number of objects and thus ensure a statistically small error for ***p***(***f***). The resulting ***p***(***f***) values were fit to a Gaussian distribution (Figure 1D), resulting in mean ***μ_fit_*** and standard deviation ***σ_fit_***. As further validation, we found ***σ_fit_*** to correspond well to the core diameter setting in the INSPIRE software (Figure S6).

### Multicolor channel registration

To compare localizations made between different spectral channels, *i.e*. different parts of the detector, it is necessary to perform subpixel, cross-channel registration. Due to field-dependent aberrations that might affect the channels differently, we divided the imaging volume into 8×8×8 voxels and corrected for the average displacement between two channels. This data was then interpolated in 3D, to create a function that corrects each localization by the estimated bias at that point in the field, *X’Y’Z’* = *f*(*X, Y, Z*) (Figure S4).

### Removing outlier localizations and low SNR objects

For calibration, fluorescent objects were required to be localizable in each channel used to determine the z distribution, and had to have an amplitude at least 3X larger than the standard deviation of the background.

For *in vitro* DNA experiments, objects with a measured 3D distances >0.5 μm, comprising <2.7% of the total analyzed objects measured in flow and <0.3% of objects on the coverslip, and were excluded when calculating the mean and standard deviations of the distributions.

For cell measurements, a minimum amplitude of the Gaussian fit was required to ensure that the spot was localizable [11] D.U. and chosen because the distribution broadened at lower thresholds but did not change significantly at higher thresholds.

### Simulation of the Tetrapod point-spread function

To simulate the Tetrapod point spread function a numerical model of the imaging system was constructed based on a scalar approximation model as described previously (Shechtman et al., 2016). The simulated emission path of the ISX contained an objective lens (60X, NA 0.9), a tube lens, and relay lenses which form the diffraction-limited image on the detector. The phase mask was inserted in a conjugate back focal plane to the objective. The physical phase mask was simulated as a phase-modulating element combined with an iris described by:

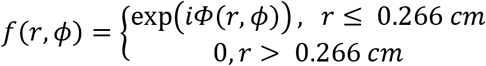

Where *r, ϕ* are the polar coordinates in the back focal plane, and Φ is the phase mask design (Figure 6 C) described previously (Shechtman et al., 2015). Due to the refractive index mismatch between the sample medium (approximately equal to that of water, *n* = 1.33) and immersion medium of the objective (air, *n* = 1), a depth change of the emitter manifests as a stronger defocus in phase than the equidistant change in objective position. The distance of the objective relative to the flow cell was therefore determined empirically.

### Data availability

The ImageStream data (.cif and .txt files) will be made available upon request.

## APPENDIX AND SUPPLEMENTARY FIGURES

### Signal-dependent precision

To evaluate the localization precision at a variety of signal levels, fluorescent beads were imaged at varying laser power [30,200] mW. The precision between channels 2 and 4 were then evaluated as a function of the localized amplitude (Figure S5).

### Preparation and imaging of DNA nanorulers by standard microscopy

For standard microscopy, nanorulers were prepared using the protocol provided by the manufacturer (Schmied et al., 2014). Briefly, clean coverslips with a custom PDMS (polydimethylsiloxane) chamber were washed three times by pipetting 400 μl of PBS (phosphate buffered saline, Sigma P5368; 1L contains NaCl 0.138 M, KCl 0.0027 M; pH 7.4). Coverslips were then incubated for 5 min with 200 μl BSA-biotin (biotin-labeled bovine serum albumin, Sigma cat. A8549) diluted to 1mg/1ml in PBS (initial dilution of 1 mg in 900 μl ddH_2_O [doubly-distilled water] followed by addition of 100 μl 10X PBS). BSA-biotin solution was removed by pipetting and coverslips were washed three times with 400 μl PBS. Coverslips were incubated for 5 min with neutravidin (Sigma cat. 31000) diluted to 1 mg/ml in PBS (initial dilution of 1 mg in 900 μl ddH_2_O followed by addition of 100 μl 10xPBS). Neutravidin solution was removed and coverslips were washed three times with 400 μl PBS/10mM MgCl2 (stock solution of 1M MgCl_2_ was prepared from anhydrous MgCl_2_, Alfa Aesar cat. 12315). 1ul of DNA nanoruler sample (stock concentration 2-4 nM) was diluted in PBS/10mM MgCl_2_. The entire 200 μl were then deposited on the coverslips. All liquid components were vortexed briefly before deposition. The deposition protocol was carried out at room temperature.

Radial, 2D distances between the fluorescently-labeled ends of the DNA nanorulers were imaged using an inverted Nikon Ti microscope outfitted with a 100X, NA 1.45 objective lens (also Nikon) and SCMOS detector (Prime 95B, Photometrics). Imagedata was first analyzed using the Fiji distribution of ImageJ (Schindelin et al., 2012) with the Thunderstorm plugin (Ovesný et al., 2014). Color-channel registration of images was done using an affine transform. Example images and histograms are shown in Figure S2.

### Calibration-object-dependent precision

The calibration procedure developed in this work relies on the convergence of a sample distribution to its mean as illustrated in Figure S1. Experiments in this work utilize at least 10000 calibration objects to form the shape-to-depth curve, corresponding to a depth error of ^~^30 nm for each calibration objects in the typical flow-core conditions (7 μm diameter), absent of localization precision and other errors described.

### Supplemental figures

**Figure S1.**
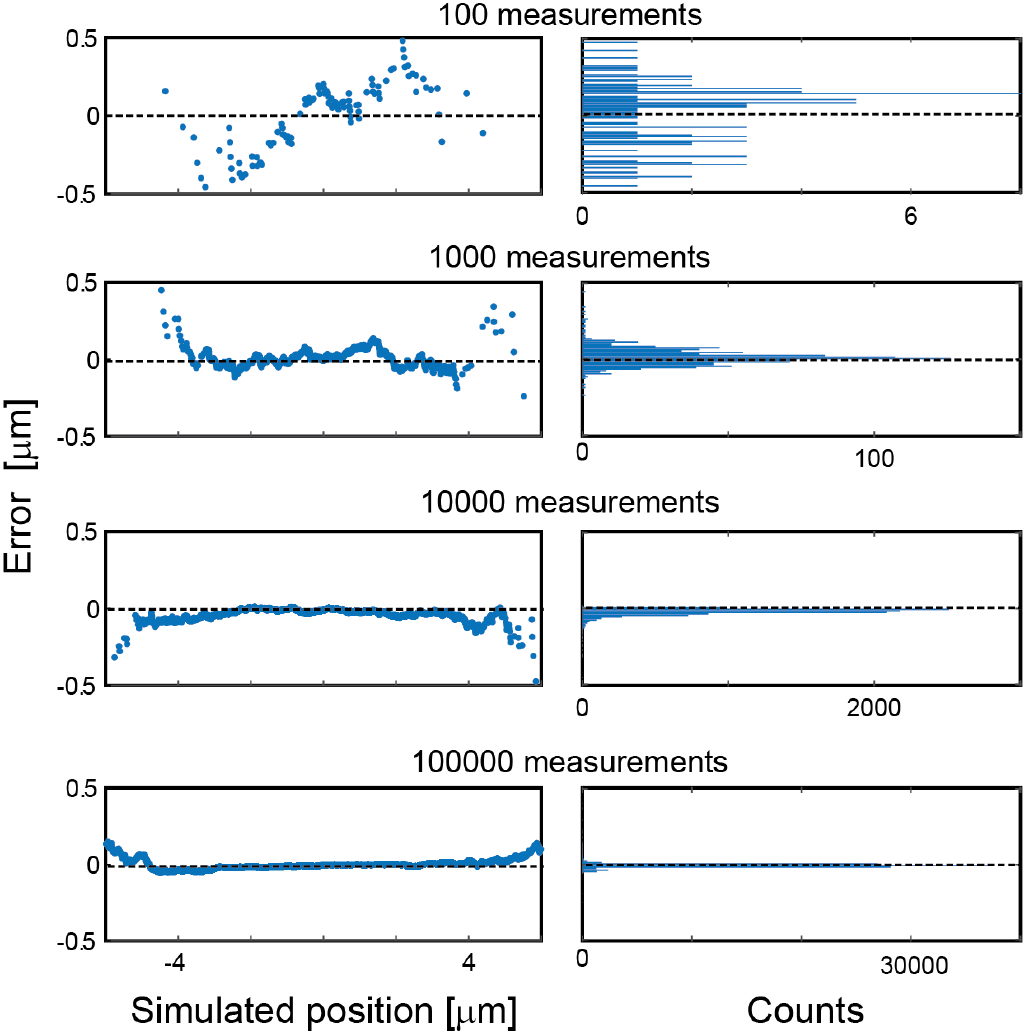
Simulated calibration test. A set of N simulated calibration object measurements were drawn from a normal distribution with mean 0 μm and standard deviation of 1.75 μm. The error was then defined as the difference between the simulated positions to the statistically most likely position for a given number of measurements. Note that this calculation does not take into account the localization uncertainty of an individual measurement.

**Figure S2.**
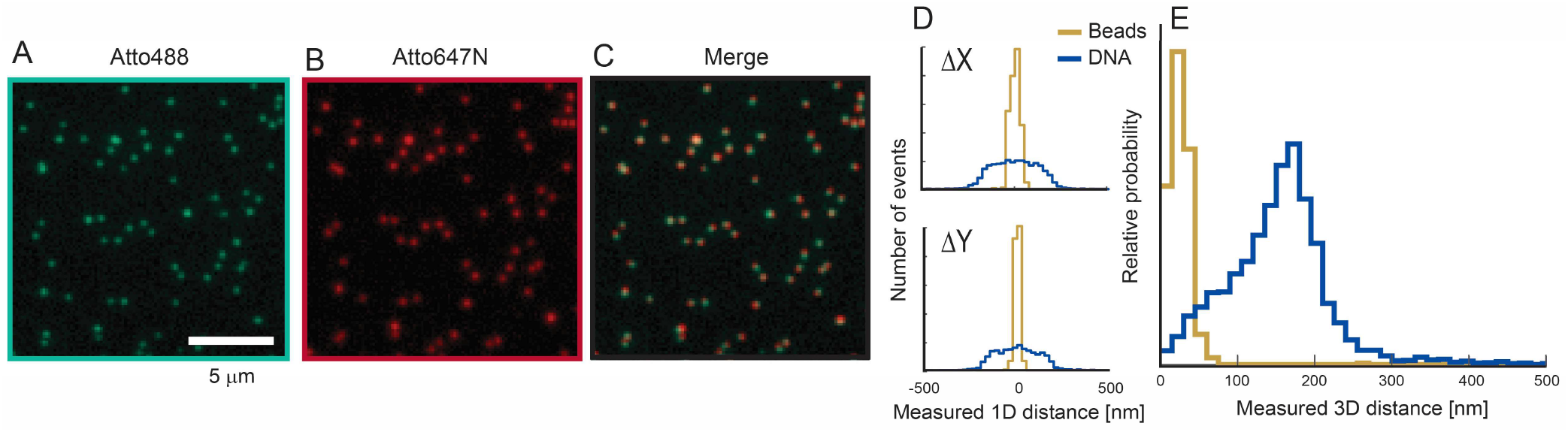
Nanorulers measured by standard microscopy. Fluorescent nanorulers were fixed to a glass slide with biotin and imaged sequentially (A) Atto488 (B) Atto47N, then aligned via an affine transformation using multicolor beads. (D) 1D and (E) radial displacements.

**Figure S3.**
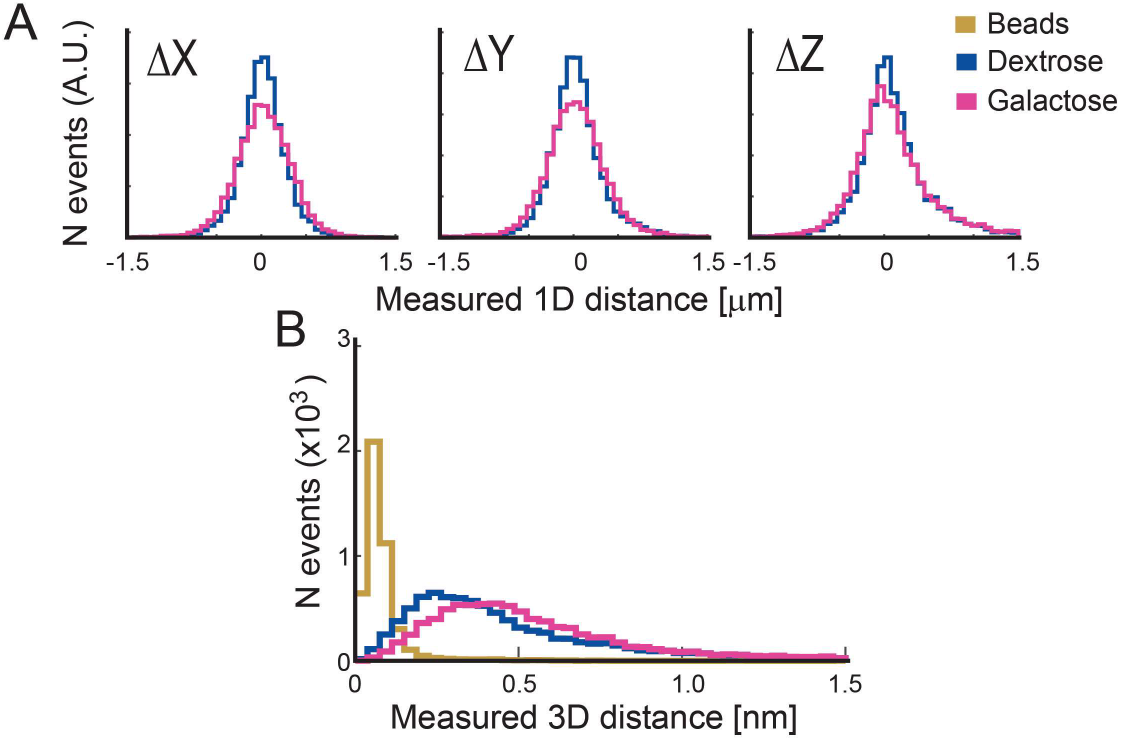
3D compaction of the Gal locus in fixed cells. (A) Histogram of inter-channel distance between localizations in each dimension and (B) 3D. Mean ± standard deviations 529 ± 286 and 455 ± 293 for galactose (N = 8432) and dextrose (N = 8646), respectively.

**Figure S4.**
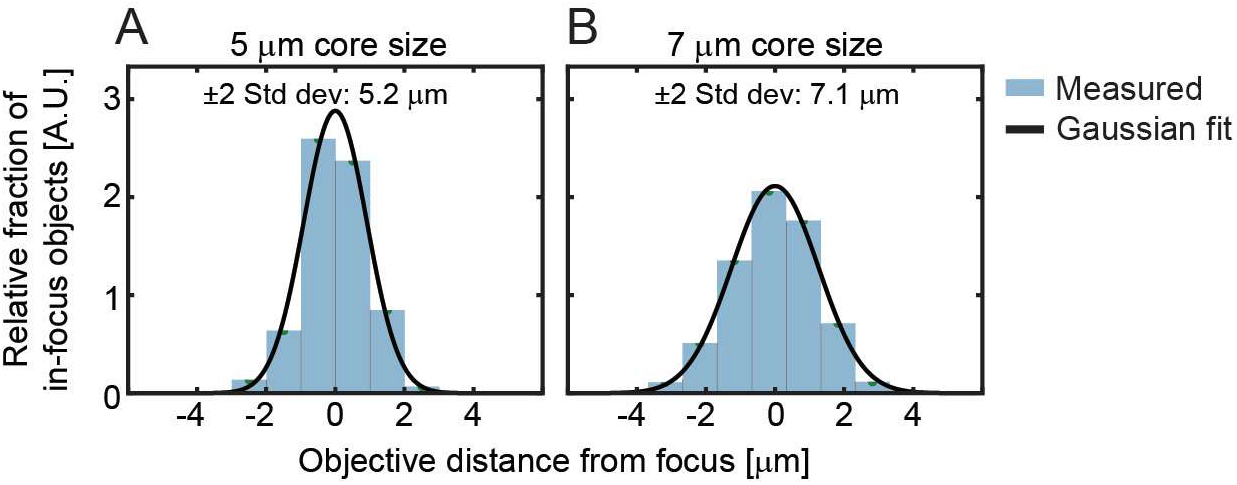
Z distributions of objects measured for two flow width parameter settings (A) 5 μm and (B) 7 μm.

**Figure S5.**
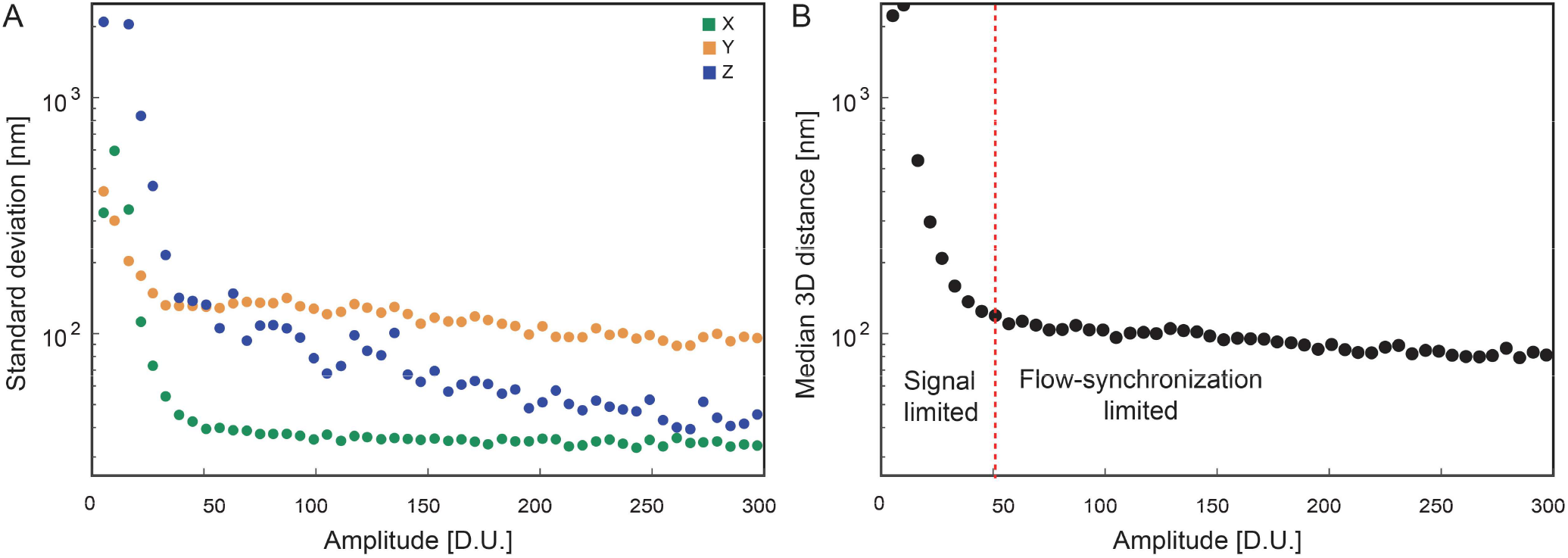
3D precision as a function of amplitude for fluorescent beads. (A) the standard deviation in X, Y, & Z as a function of the localized amplitude. (B) The median 3D distance over two regimes: signal limited and flow-synchronization limited.

**Figure S6.**
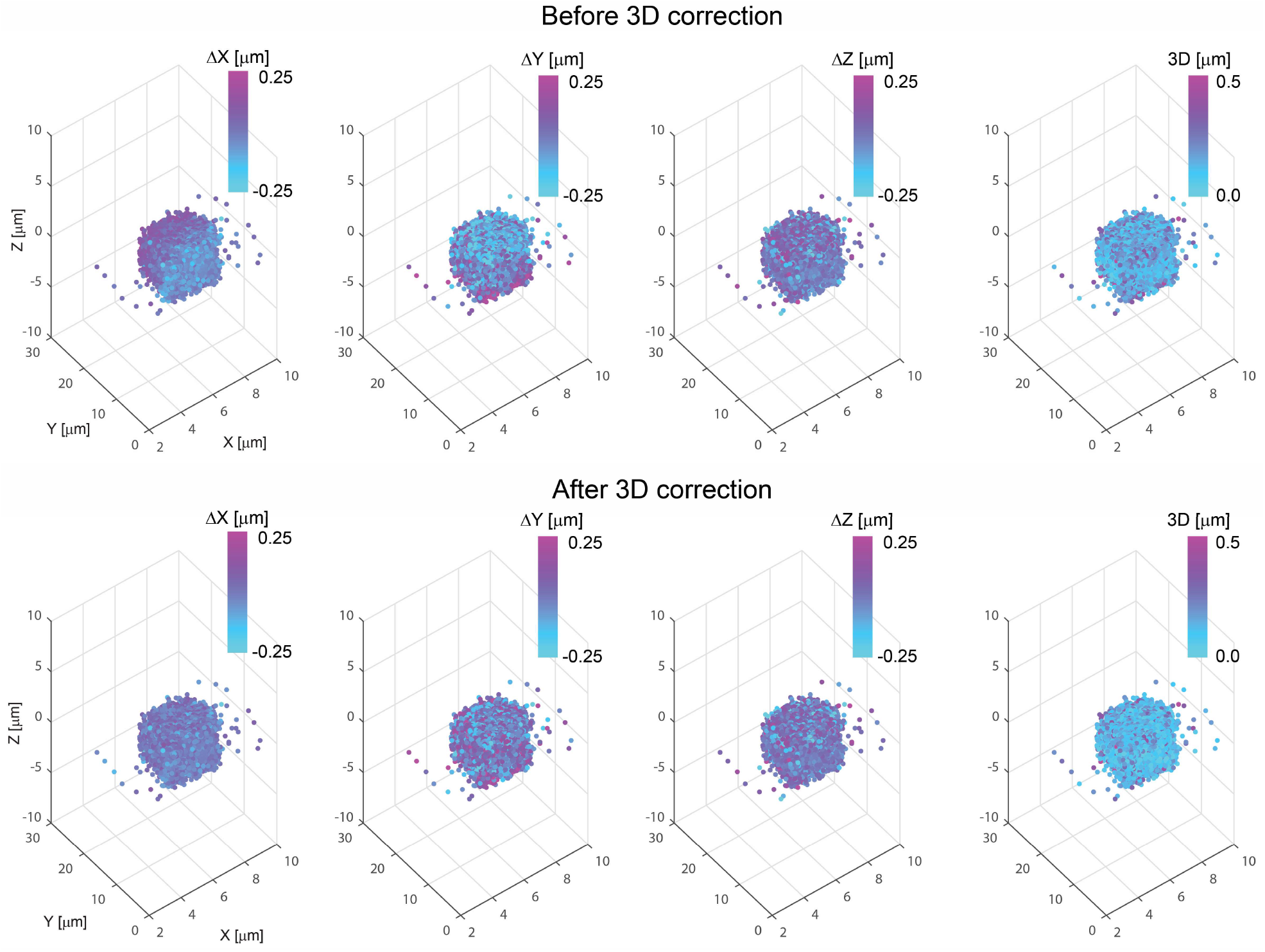
3D correction of localization bias. (A) Bead localizations (A) before and (B) after 3D corrections. Each point represents a single bead colored by the displacement in each axis and in 3D (right).

